# Development of a Virus Particle-Based Antibody-Dependent Cellular Phagocytosis (ADCP) Assay for HIV

**DOI:** 10.1101/2025.08.27.672700

**Authors:** Chitra Upadhyay, Priyanka Gadam Rao

## Abstract

Antibody-dependent cellular phagocytosis (ADCP) is an Fc-mediated effector function that contributes to the clearance of pathogens, including HIV. Conventional bead-based ADCP assays using recombinant Env proteins offer scalability and reproducibility but do not recapitulate the structural, conformational and glycan features of Env on intact virions, potentially limiting their physiological relevance. Here, we describe a virus particle-based ADCP assay that preserves the native membrane embedded conformation and glycosylation of the HIV envelope glycoprotein (Env), features critical for antibody recognition and Fcγ receptor engagement. The assay utilizes THP-1 human monocytic cell line, as effector cells and sucrose purified inactivated HIV-1 virions coupled with fluorescent beads, as phagocytic targets. This platform enables sensitive and reproducible measurement of antibody-mediated phagocytosis in a biologically relevant antigen presentation context. The assay supports functional profiling of monoclonal and polyclonal antibodies across multiple species and provides a practical framework for evaluating Fc effector responses in HIV infection and vaccine studies.

## Introduction

Broadly neutralizing antibodies (bNAbs) are a major focus of HIV vaccine and therapeutic research due to their demonstrated ability to prevent infection and suppress viraemia in both preclinical and clinical settings. In nonhuman primate (NHP) and humanized mouse models, passive administration of bNAbs has been shown to protect against simian-human immunodeficiency virus (SHIV) and HIV challenge respectively (1–7). Clinical trials, including the Antibody Mediated Prevention (AMP), have further validated this approach, demonstrating up to 75% protection against bNAb-sensitive HIV-1 strains in humans (8). These findings highlight the central role of bNAbs as correlates of protection.

The predominant mechanism underlying bNAb-mediated protection is Fab-mediated neutralization, where antibodies block viral entry by targeting the conserved regions of the HIV-1 envelope glycoprotein (Env) (9, 10). However, accumulating evidence indicates that Fc-mediated effector functions such as antibody-dependent cellular phagocytosis (ADCP), antibody-dependent cellular cytotoxicity (ADCC), and complement activation, also contribute to antiviral efficacy, facilitating clearance of infected cells and virions (11–18). The interplay between Fab- and Fc- dependent mechanisms is increasingly recognized as critical for the breadth and durability of bNAb-mediated protection.

In the context of HIV-1, ADCP has been associated with reduced infection risk in NHP vaccine studies (19, 20). In human clinical trials, including RV144 and HVTN 505, Fc mediated antibody functions, including ADCP activity, were detected as part of the vaccine elicited humoral response (21). These findings highlight the need to better understand and quantify ADCP as a correlate of protective immunity. Several assay platforms have been developed to evaluate ADCP in the context of HIV. The most widely adopted approach utilizes beads coated with HIV Env proteins, which are opsonized by antibodies and subsequently internalized by monocytic cell lines such as THP-1 (22–27). This bead-based assay format is highly scalable and reproducible, enabling quantitative assessment of Fc-mediated phagocytic activity across diverse antibody specificities. Infected cell-based assays offer greater physiological relevance by presenting native trimeric Env on the surface of infected cells targets (28), but they are technically demanding and subject to variability in antigen density and cell viability (29–31). Importantly, while these platforms differ in complexity and scalability both formats share a key limitation: they do not enable assessment of Fc-mediated phagocytosis in the context of authentic viral particle-associated Env (31, 32). Conventional bead-based systems using recombinant Env proteins (e.g., gp120, gp140, V3 or V1V2 proteins) (24–27) oversimplify Env presentation, omitting critical conformational and glycan features of the native trimer, and thus do not fully recapitulate the antigenic landscape of intact virions. Infected cell assays incorporate viral antigens in a more native cellular context but do not directly interrogate particle-level constraints on antibody-mediated clearance and are less amenable to high-throughput or cross-study standardization. Collectively, these limitations restrict our ability to evaluate how antigen context and Fc effector function cooperate to shape antibody-mediated phagocytic activity.

The aim of this study was to develop a virion-based ADCP assay that enables direct measurement of antibody-mediated phagocytosis of HIV virions. By coupling native HIV virions to fluorescent beads, this approach integrates the biological relevance of viral particles with the scalability and reproducibility of conventional bead-based platforms. The combination of physiologically relevant antigen presentation and high-throughput compatibility makes the virion-bead ADCP assay well-suited for vaccine evaluation, systems serology, and translational studies focused on identifying correlates of protective immunity. This platform supports robust functional profiling in both research and clinical settings.

## Materials and Methods

### Cell lines

The HEK293T/17 (293T) were obtained from the American Type Culture Collection (ATCC, Manassas, VA). The following reagent was obtained through the NIH HIV Reagent Program, Division of AIDS, NIAID, NIH: TZM-bl Cells, ARP-8129, contributed by Dr. John C. Kappes, Dr. Xiaoyun Wu, and Tranzyme Inc (33). For all experiments, HEK293T/17 cells (293T) were used to produce infectious HIV-1 viruses and the TZM.bl cell line was used to assay virus infectivity. TZM.bl cell line is derived from HeLa cells and is genetically modified to express high levels of CD4, CCR5 and CXCR4 and contains reporter cassettes of luciferase and β-galactosidase that are each expressed from an HIV-1 LTR. The 293T and TZM.bl cell lines were routinely sub-cultured every 3 to 4 days by trypsinization and were maintained in Dulbecco’s Modified Eagle’s Medium (DMEM) supplemented with 10% heat-inactivated fetal bovine serum (FBS), HEPES pH 7.4 (10 mM), L-glutamine (2 mM), penicillin (100 U/ml), and streptomycin (100 μg/ml) at 37°C in a humidified atmosphere with 5% CO2.

HEK293F cells (Thermo Fisher Scientific) were used to produce mouse mAbs in this study. The cells were cultured and maintained according to the manufacturer’s recommended protocols.

THP-1 cells are a human monocytic cell line commonly used in ADCP assays to model macrophage-mediated phagocytosis. THP-1 cells were purchased from ATCC and maintained in RPMI 1640 media (ATCC) containing 2 mM L-Glutamine (Gibco), 10% Fetal Bovine Serum (Sigma), 10 mM HEPES (Gibco), 55 μM beta-mercaptoethanol (Gibco), and 1X Penicillin/Streptomycin (Gibco). Cell culture densities were kept below 0.5 × 10^6^ cells/ml to maintain consistent assay performance.

### Plasmids

A full-length transmitted/founder (T/F) infectious molecular clone (IMC) of pREJO.c/2864 (REJO, ARP-11746) was obtained through the NIH HIV Reagent Program, Division of AIDS, NIAID, NIH, contributed by Dr. John Kappes and Dr. Christina Ochsenbauer (34). REJO is a tier 2, clade B, T/F isolate. The IMC of RHPA and QH0692 (tier 2, clade B, T/F) were kindly provided by Dr Benjamin Chen. The IMCs were generated by cloning the Env into a pNL4.3 backbone to construct pNL-RHPA and pNL-QH0692, respectively (35). The CMU06 IMC was generated similarly by cloning the Env into a pNL4.3 backbone to construct pNL-CMU06(35).

### Antibodies and plasma samples

The following antibody reagents used in this study were obtained through the NIH AIDS Reagent Program, Division of AIDS, NIAID, NIH: Anti-HIV-1 gp120 monoclonal VRC01 from Dr. John Mascola (36); anti-HIV-1 gp120 monoclonal antibody NIH45-46 G54W, contributed by Dr. Pamela Bjorkman (37); anti-HIV-1 gp41 monoclonal 2F5 from Polymun Scientific (38); polyclonal HIV-Ig contributed by Dr. Luiz Barbosa. The V2i (830) and V3 (2219) and the irrelevant (non-HIV-1) anti-anthrax 3685 mAbs were obtained from the laboratory of Dr. Susan Zolla-Pazner (39–47). The mAb 3685 was used as a negative control. HIV-1 positive human plasma samples used were a gift from Dr Colleen Courtney (48). The mouse serum samples were from animals co-immunized with gp120 DNA and protein immunogens (27). Mouse monoclonal antibodies were generated by hybridoma technology, where splenic B cells of mouse immunized with gp120 immunogens were fused with myeloma cells to create antibody-secreting hybridomas. For production and purification, the antibody sequences (heavy and light chains) were cloned into the mammalian expression vector pcDNA3.1. These plasmids were then co-transfected into HEK293F cells for protein expression (data not shown). The transfected cells expressed the recombinant antibodies, which were subsequently harvested from the culture supernatant, purified and used as reagents in this study.

### Virus production and purification

Infectious viruses were generated by transfecting 293T cells with pREJO, pNL-CMU06, pNL-RHPA and pNL-QH0692 plasmids using jetPEI transfection reagent (Polyplus, New York, NY) (49). Supernatants were harvested after 48 hours and clarified by centrifugation and 0.45μm filtration. Virus infectivity was assessed on TZM.bl cells as described(35, 49). Briefly, serial two-fold dilutions of virus stock in 10% DMEM were incubated with TZM.bl cells (in duplicates for each dilution) in half-area 96-well plates in the presence of DEAE-dextran (12.5 μg/ml) for 48 hours at 37°C. Virus infectivity was measured by β-galactosidase activity (Promega, Madison, WI). Virus stocks were concentrated (20X) by ultracentrifugation over 20% (w/v) sucrose in 1X phosphate buffered saline (PBS) at 25,000 RPM for 2 hours in an SW-28 swinging bucket rotor (Sorvall, Thermofisher Scientific). Supernatants were decanted and pellets dried briefly before resuspension in PBS. Inactivation of virions was carried out using Aldrithiol-2 (AT-2) (50, 51). Briefly, 125 ul of sucrose-purified virus was incubated with 0.5 mM AT-2 in DMSO for 2 hours at 37^°^C, followed by centrifugation at 13,000 rpm for 2 hours. The supernatant was discarded, and the pellet re-suspended in 125 ul PBS. Inactivation was confirmed by measuring infectivity in TZM.bl cells and Env content was checked by Western blotting.

### Western blotting

To quantify and monitor the expression of Env in each virus preparation Western blot analyses were performed as described before (52). Briefly, the sucrose-purified virus particles were lysed, resolved by SDS-PAGE on 4-20% tris-glycine gels (Bio-Rad, Hercules, CA), and blotted onto membranes, which were then probed with antibodies. A cocktail of anti-human anti-gp120 MAbs (anti-V3: 391, 694, 2219, 2558; anti-C2: 841, 1006; anti-C5: 450, 670, 722; 1μg/ml each) was used to detect Env. MAb 91-5D (1μg/ml) was used to detect Gag p24. Membranes were developed with Clarity Western ECL Substrate (Bio-Rad, Hercules, CA) and imaged by iBright FL1500 Imaging System (Invitrogen, Carlsbad, CA). Purified recombinant gp120 and p24 proteins were also loaded at a known concentration as controls and quantification standards. Band intensities were quantified using the iBright Analysis Software Version 5.0.1 (Invitrogen, Carlsbad, CA).

### Coupling of fluorescent beads to virions

The coupling strategy used in this study is a standard method widely used in Luminex based multiplex immunoassays and is published previously (53–56). Here we applied this approach for coupling the virus to the beads (52). Sucrose-purified, inactivated virions were covalently coupled to carboxylate-modified FluoSphere microspheres (1.0 µm) using a two-step carbodiimide reaction with the xMAP Ab Coupling (AbC) Kit according to manufacturers’ instructions (Luminex, Austin, TX). Carboxylated beads purchased from Thermo Fisher (cat# F8823; 505/515 nm), were coupled to 125 μl of 20X concentrated virus preparations. Briefly, the stock microspheres were vortexed and sonicated to resuspend and disperse the microspheres and 12 μl (∼36.4x10^9^ beads) was transferred to a tube containing 1200 ul of 1% BSA/1X PBS (per virus). The microspheres were washed twice with 500 μl of activation buffer followed by vortexing and sonication after each step. The microspheres were activated with 400 μL of activation buffer, 50 μL of 50 mg/ml Sulfo-NHS (N-hydroxysulfosuccinimide), 50 μL of 40 mg/mL ethyl dimethylaminopropyl carbodiimide hydrochloride (EDC) and incubated for 20 min at room temperature with end-to-end rotation. The microspheres were washed three times in activation buffer, then incubated with AT-2 inactivated virus in activation buffer for 2 hours at room temperature. We typically used a volume of 20X concentrated virus that equals ∼175ng total as measured by western blotting. The microspheres were subsequently washed and resuspended in 1.2 mL of 0.1% BSA/PBS and stored at 4°C until ready to use. This volume is sufficient to run the assay on two 96 well plates.

Based on the theoretically estimated Env content of HIV-1 virions (∼10-14 Env trimers per particle, corresponding to ∼30-40 gp120 molecules per virion), coupling 175 ng of virion-associated Env to approximately 3.6 × 10¹ beads corresponds to a theoretical maximum of ∼2-3×10¹ virions per reaction, or on the order of one virion per bead. Because this estimate assumes complete recovery and 100% coupling efficiency, the actual number of virions per bead is likely lower. Based on these theoretical considerations, the bead-to-virus ratio was selected to minimize the likelihood of multiple virions conjugating to a single bead and to favor low-density Env presentation that more closely approximates native viral conditions.

### ADCP Assay Protocol

The coupled microspheres (10 µl/well) were incubated with serial dilutions of monoclonal antibodies or plasma (10 µl/well) for 2 hours at 37°C. Antibody opsonized virions coupled microspheres were washed with 0.1% BSA/PBS twice to remove unbound antibodies. THP-1 cells were added (200 μL/well) at a concentration of 1.25 × 10^5^ cells/mL (2.5 × 10^4^ cells/well) and incubated with the immune complexed microspheres overnight for ∼16 h at 37°C. Cells were then fixed with 4% PFA and acquired on a Attune NxT flow cytometer.

Data analysis was performed using FCS Express 7 Research Edition (De Novo Software) as follows: Cells were gated on a plot of forward scatter area versus side scatter area (FSC-A vs SSC-A). Doublets were excluded using a forward scatter height versus area plot (FSC-H vs FSC-A). Geometric mean fluorescence intensity (MFI) values of AF488 cells, representing phagocytosed virus particles coupled to yellow green microspheres, were determined. ADCP scores were calculated as follows: [(% microsphere positive cells) x (MFI of the microsphere positive cells)/10,000]. Control wells including virion coupled beads alone, cells alone and virion coupled beads without Ab and cells (no Ab control for background phagocytosis).

### Statistical analysis

Statistical analyses were performed as indicated in the figure legends with one or two-way ANOVA using Dunnett multiple comparison test using GraphPad Prism 10.6.0. Statistical significance was interpreted in conjunction with effect size and dose-response behavior to avoid overinterpretation of small but consistent differences.

Intra-assay precision was calculated as the coefficient of variation (CV%) between technical replicates within a single experiment, and inter-assay reproducibility was calculated as the CV across independent experiments performed on different dates using the mean replicate value per run. Mouse mAbs tested in Fig 5 with RHPA virions were used.

## Results

### Development of a virus particle-based ADCP assay

To establish a biologically relevant platform for measuring Fc effector activity against HIV-1, we developed a virus particle-based antibody-dependent cellular phagocytosis (ADCP) assay. As shown in Figure 1A, sucrose-pelleted RHPA virions were coupled to 1 μm yellow-green, fluorescent beads via a two-step carboxyl chemistry. Bead coupling was used to generate a phagocytosis-competent target while preserving native, membrane-associated Env presentation on intact viral particles. These virion-coated beads were opsonized with antibody and incubated with THP-1 monocytic effector cells. After fixation, phagocytosis was quantified by flow cytometry using AF488 fluorescence to detect bead uptake by THP-1 cells. Gating was performed on singlet THP-1 cells to exclude aggregates, and ADCP scores were calculated by multiplying the geometric mean fluorescence intensity (MFI) of bead-positive cells by the percentage of bead-positive cells, normalized to background.

**Figure 1.**
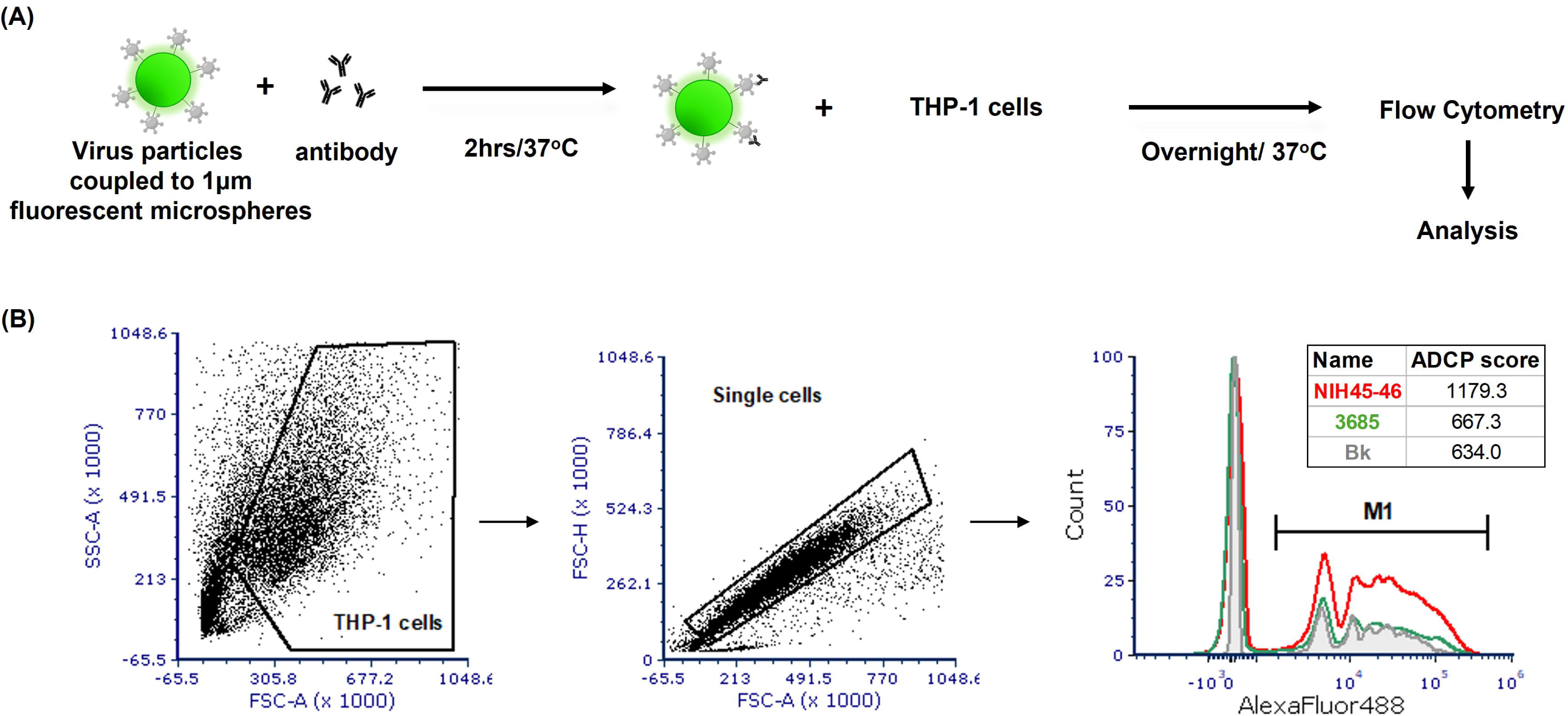
ADCP Assay for Detecting Phagocytosis of Antibody-Opsonized HIV-1 Virions. **(A)** Schematic overview of the assay. Sucrose pelleted RHPA virions were carboxyl-coupled to 1 μm yellow-green beads. Virion-coupled bead were then incubated with HIV-1 envelope-specific mAb, washed, incubated with human THP-1 monocytes, fixed, and analyzed by flow cytometry. **(B)** Gates were drawn on singlet THP-1 cells, and phagocytic scores were calculated from data on the AlexaFluor488 fluorescence channel to quantify envelope-specific ADCP. Histograms indicate results with 5 μg/ml positive (NIH45-46; red) and negative control (3685-non-HIV-1; green) mAbs. Background phagocytosis (Bk) of virion-couple beads with no Ab is shown in gray.

In our previously published study (52), we optimized the amount of Env used for bead coupling to achieve a low signal-to-noise ratio while maintaining clear separation between positive and negative antibody binding. For the ADCP assay reported here, we used the same optimized Env amount for conjugating to the beads. Initially, we used RHPA, a T/F, clade B, tier 2, HIV-1 isolate and monoclonal antibodies (mAb) that recognize the conformational CD4bs (NIH45-46) epitope on Env. A non-HIV-1 mAb 3685 was used as a negative control. As shown in Figure 1B, the mAb NIH45-46 (red histogram) mediated robust phagocytic activity with an ADCP score of 1179.3, which was higher than both the non-HIV-1 control mAb 3685 (green histogram; ADCP score = 667.3) and the background control with no antibody (gray histogram; ADCP score = 634.0). These results confirm the specificity and sensitivity of the assay for detecting Fc-mediated phagocytosis of antibody-opsonized HIV-1 virions.

### Detection of ADCP activity by human monoclonal antibodies

We next applied the assay to evaluate the functional capacity of a panel of well-characterized human HIV-1 mAbs targeting diverse HIV-1 Env epitopes including the CD4 binding site (VRC01, NIH45-46), gp41 (2F5), V2 (830), and V3 linear epitopes (2219). Each mAb was tested across a range of concentrations using RHPA HIV-1 virion coupled to fluorescent beads and THP-1 monocytes. As shown in Figure 2A-B, some HIV-1-specific mAbs mediated dose-dependent phagocytosis, with NIH45-46 and 830 demonstrating the highest ADCP activity, followed by moderate responses from VRC01 and 2F5 (Fig 2A-C). As expected, the irrelevant anthrax-specific mAb 3685 showed no detectable activity, confirming assay specificity. The V3 mAb 2219 did not mediate detectable ADCP. Quantitative analysis of the area under the titration curves (AUC) from both experiments (Figure 2C) confirmed significant differences in potency across the panel, with NIH45-46 and 830 yielding the highest values. The irrelevant anthrax-specific mAb 3685 remained at baseline throughout, validating assay specificity.

**Figure 2.**
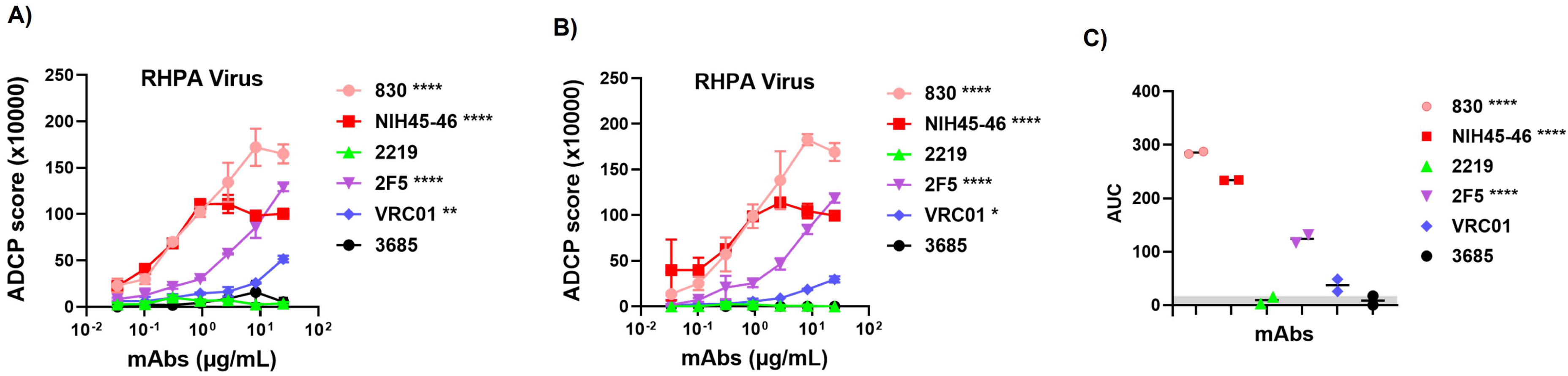
ADCP assay for detection of phagocytosis of human antibody opsonized virions. Sucrose pelleted RHPA virions were carboxyl-coupled to 1 μm yellow-green beads. Virion-coupled bead were then incubated with HIV-1 envelope-specific mAbs, washed, incubated with human THP-1 monocytes, fixed, and analyzed by flow cytometry. Beads incubated with cells without Ab were used as background and subtracted. An irrelevant non-HIV-1 mAb specific to anthrax (3685) was used as negative control. **(A)** Experiment 1 and **(B)** Experiment 2 showing dose-response curves of ADCP activity of HIV-1 mAbs against RHPA virions across a 3-fold dilution series starting at 25 μg/mL. Data are shown as mean ± SD from replicate wells. **(C)** Area under titration curve (AUC) values calculated from panel A and B. Gray shading indicates the AUC for the negative control mAb (3685). Each dot represents an independent experiment, and the horizontal line indicates the median. For panels A and B, statistical significance was determined using a two-way ANOVA with Dunnett’s multiple-comparisons test, comparing each mAb to the negative control (3685) based on the main column effect averaged across all concentrations. Significance levels are indicated as *, p<0.05; **, p<0.01; *** p<0.001; ****, p<0.0001. Values with p > 0.05 are unmarked. For panel C, a one-way ANOVA with Dunnett’s multiple-comparisons test was used to compare the AUC of each mAb to the negative control (3685).

### Broad applicability across viral strains and antigens

To assess the applicability of the virion-based ADCP assay across diverse HIV-1 strains and antigen formats, we tested the same mAb panel against virions from two additional isolates, CMU06 and REJO as well as recombinant REJO gp120. With CMU06 virions (Figure 3A), mAbs 830 and 2219 mediated robust, dose-dependent ADCP responses across the concentrations tested. Among all antibodies, 830 and 2219 displayed the strongest phagocytic activity, with ADCP scores markedly exceeding those of all other mAbs at mid- to high concentrations. NIH45-46 and 2F5 showed moderate activity above background, whereas VRC01 remained near baseline, similar to the irrelevant control antibody 3685. When tested against REJO virions (Figure 3B), mAb 830 retained strong dose-dependent phagocytic activity, while NIH45-46 and 2F5 showed only low activity. V3 mAb 2219, which was effective against CMU06, exhibited low activity and did not mediate appreciable ADCP. These findings highlight strain-dependent differences in antibody-mediated phagocytosis, consistent with variation in Env epitope accessibility between viral isolates (52) and highlights the importance of testing multiple HIV-1 strains rather than relying on a single isolate.

**Figure 3.**
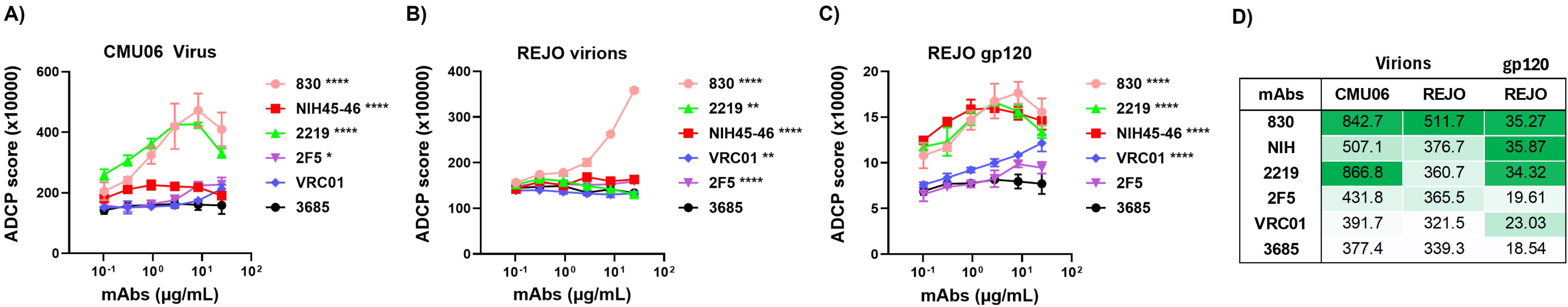
Broad Applicability of the ADCP Assay Across HIV-1 Strains and Antigen Formats. Sucrose pelleted virions were carboxyl-coupled to 1 μm yellow-green beads. Virion-coupled bead were then incubated with HIV-1 envelope-specific mAb,s washed, incubated with human THP-1 monocytes, fixed, and analyzed by flow cytometry. ADCP with **(A)** CMU06 virions, **(B)** REJO virions and **(C)** Rejo gp120. Beads incubated with cells without Ab were used as background and subtracted. An irrelevant non-HIV-1 mAb specific to anthrax (3685) was used as negative control. Data represent the mean ± SD of replicate measurements from one experiment. (D) Area under the curve (AUC) values were calculated from panel A, B and C. Color-code is relative to the AUC of the negative control mAb (3685) for each antigen format, with darker green indicating larger effect sizes compared to 3685. Higher AUC values reflect stronger phagocytic activity. Statistical significance was determined using two-way ANOVA with Dunnett’s multiple-comparisons test comparing each mAb to the negative control (3685) based on the main column effect averaged across concentrations (*p < 0.05; **p < 0.01; ***p < 0.001; ****p < 0.0001). Values with p > 0.05 are unmarked.

We next assessed ADCP activity against recombinant Env by testing the panel with REJO gp120 monomers coupled to beads (Figure 3C). In this format, mAbs 830, NIH45-46, and 2219 mediated the strongest phagocytic responses. VRC01 consistently showed weak activity with both gp120 and virions. As expected, the gp41-specific mAb 2F5 did not show any response. Because antigen density and presentation differ substantially between virion and protein coupled beads, direct quantitative comparison of ADCP magnitude across antigen formats were not performed. As anticipated, the gp120-based assay did not replicate the restricted ADCP pattern seen with REJO virions. Instead, antibodies that were weak against REJO virions, such as 2219 and NIH45-46, mediated strong phagocytic responses with gp120-coupled beads, highlighting the impact of Env organization and membrane context on epitope accessibility and Fc-effector function.

To summarize differences in ADCP magnitude within each antigen format, we calculated the area under the curve (AUC) for each mAb. Figure 3D presents a color-coded comparison of AUC values compared to the negative control (mAb 3685), providing an visual summary of antibody-mediated phagocytosis across antigen contexts. For virions, mAb 830 exhibited the largest enhancement over 3685 for both CMU06 and REJO (AUC = 842.7 and 511.7, respectively), whereas 2219 showed strong activity against CMU06 (AUC = 866.8) but minimal activity against REJO (AUC = 360.7). For gp120, AUC values ranged from 18.5 (3685) to 35.9 (NIH45-46). Within each antigen format, the AUC values allowed direct comparison among all mAbs. For CMU06 virions, 2219 and 830 showed the highest AUC values, followed by NIH45 46, 2F5, VRC01, and 3685. For REJO virions, 830 had the highest AUC, while NIH45 46, 2F5, 2219, VRC01, and 3685 formed a lower activity group with smaller differences among them. For REJO gp120, NIH45 46, 830, and 2219 showed the highest AUC values, whereas VRC01 and 2F5 exhibited lower activity approaching background levels defined by 3685. Using two-way ANOVA with Dunnett’s multiple-comparisons test, all HIV-1 specific mAbs showed statistically significant increases in mean ADCP activity relative to the negative control (3685) when averaged across concentrations. However, statistical significance did not uniformly correspond to large biological effects. Notably, the magnitude of enhancement differed substantially among antibodies, with mAb 830 exhibiting the strongest and most consistent activity, whereas other antibodies showed smaller effect sizes despite statistical significance. These findings emphasize that statistical significance alone may not fully capture biologically meaningful differences in Fc-mediated effector function, particularly when antigen presentation imposes native structural constraints.

### Polyclonal HIV-1 infected sera mediate virion phagocytosis

We next assessed whether the assay developed could detect ADCP mediated by polyclonal antibodies. RHPA virion-coupled fluorescent beads were incubated with heat-inactivated sera from HIV-1 infected individuals, as well as with purified polyclonal immunoglobulin G (HIV-Ig) followed by THP-1 monocytes as phagocytic effector cells. As shown in Figure 4A-B, sera from infected donors consistently mediated dose-dependent ADCP activity, whereas HIV-1 negative sera (Neg) showed minimal activity. Quantitative analysis of the area under the titration curves (AUC) confirmed significant higher ADCP activity in HIV-1 infected sera compared to negative control (Figure 4C). These findings demonstrate that naturally elicited polyclonal antibodies in chronic HIV-1 infection can direct Fc-mediated clearance of virions and validate the sensitivity and biological relevance of the virion-based ADCP assay for profiling humoral immunity.

**Figure 4.**
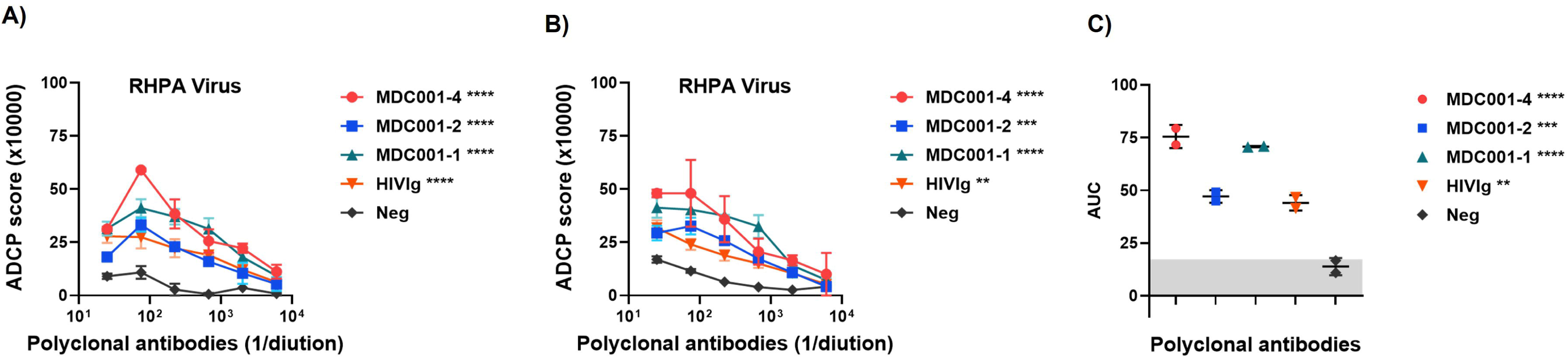
ADCP assay to measure phagocytosis of HIV-1 virions opsonized with polyclonal antibodies from infected individuals. Sucrose pelleted RHPA virions were carboxyl-coupled to 1 μm yellow-green beads. The virion-coupled bead were incubated with heat-inactivated sera from HIV-1 infected individuals and polyclonal IgG (HIV-Ig), washed, then incubated with human THP-1 monocytes. After fixation samples were analyzed by flow cytometry. Virion coupled beads incubated with cells in the absence of Ab served as background and were subtracted. Sera from healthy individual (Neg) was used as negative control. **(A)** Experiment 1 and **(B)** Experiment 2. **(C)** Area under titration curve (AUC) values were calculated from panel A and B. Gray shading indicates the AUC range for the negative control polyclonal antibody (Neg). For panels A and B, statistical significance was assessed using two-way ANOVA with Dunnett’s multiple-comparisons test, comparing each mAb to the negative control (3685) based on the main column effect averaged across concentrations. Significance is indicated *, p<0.05; **, p<0.01; *** p<0.001; ****, p<0.0001. Values with p > 0.05 are unmarked. For panel C, one-way ANOVA with Dunnett’s multiple-comparisons test was used to compare AUC values for each antibody relative to the negative control (3685). Each dot represents an independent experiment, and horizontal lines denote median values.

### Extension of the assay to mouse monoclonal antibodies and sera

To expand the utility of the ADCP assay to preclinical models, we tested murine (mu) Env-specific mAbs against RHPA and QH0692 virion-coupled beads. RHPA virions from two independent preparations (Prep 1 and Prep 2) and QH0692 virions were coupled to fluorescent beads, opsonized with mouse mAbs, and incubated with THP-1 monocytes. As shown in Figure 5A-B, mu-MAb1 mediated robust dose-dependent ADCP activity against RHPA virions, with consistent results across both virus preparations. The mu-MAb3 also mediated detectable ADCP, although with lower magnitude than mu-MAb1. In contrast, mu-MAb2 did not show any activity. The irrelevant control antibody mu-MAb4 showed no activity, and specificity was further confirmed by the Fc-impaired mu-MAb1 LALA variant, which failed to mediate ADCP (57–59). The assay also performed well with QH0692 virions (Figure 5C), where Env-specific mAbs again showed strong phagocytic responses, while mu-MAb1 LALA and mu-MAb4 remained at baseline. Notably, mu-MAb1, mu-MAb2, and mu-MAb4 are of the IgG2a isotype, whereas mu-MAb3 is IgG1. Because murine IgG2a antibodies bind activating Fcγ receptors with higher affinity than IgG1, these results are consistent with known isotype-dependent differences in Fc-mediated effector function (60). Together, these findings demonstrate that the virion-based ADCP assay is compatible with murine antibodies, reproducible across independent virus preparations, and applicable to distinct HIV-1 isolates, supporting its use for functional antibody assessment in animal models.

To further evaluate the applicability of the assay in a vaccination context, we tested serially diluted sera from mice co-immunized with HIV-1 gp120 DNA and protein were evaluated (27). Vaccinated sera mediated measurable ADCP responses in both assay formats, with variability in magnitude and dilution patterns among individual animals (Fig 6A-B). Together, these results demonstrate that the virion-coupled ADCP assay is suitable for detecting functional antibody responses induced by vaccination and complements protein-based approaches in preclinical studies.

**Figure 5.**
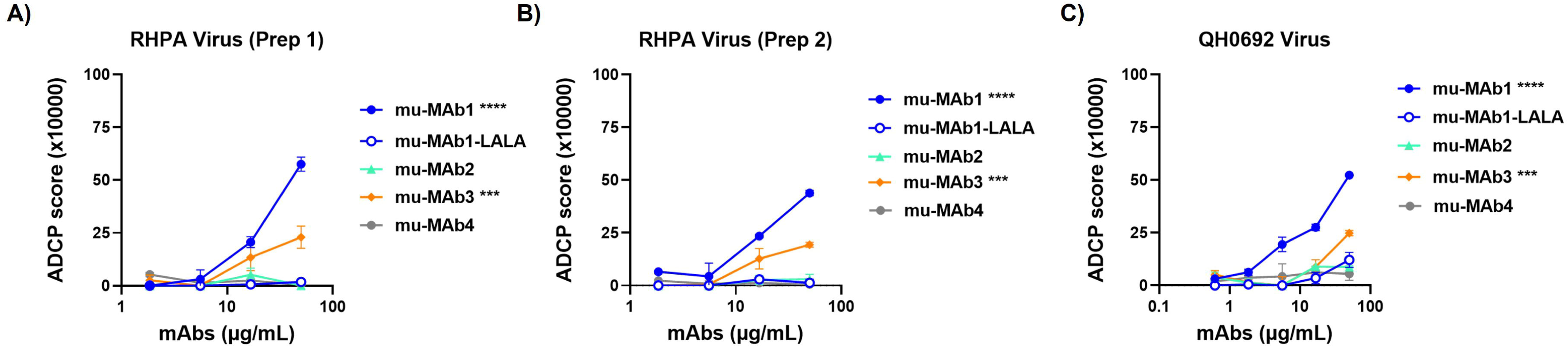
ADCP activity of murine monoclonal antibodies against HIV-1 virion-coupled beads. Sucrose pelleted RHPA virions were carboxyl-coupled to 1 μm yellow-green beads. Bead were then incubated with HIV-1 envelope-specific mouse mAb, washed, incubated with human THP-1 monocytes, fixed, and analyzed by flow cytometry. An Fc-impaired variant of mu-MAb1 (mu-MAb1-LALA) and an irrelevant murine monoclonal antibody (mu-MAb4) were included as negative controls. RHPA viruses from two different transfections are tested (**A**) Prep 1 (**B**) Prep 2). (**C**) Another T/F isolate QH0692 was tested similarly. Beads incubated with cells without Ab were used as background and subtracted. Data represent the mean ± SD of replicate measurements from one experiment. Statistical significance was determined using a two-way ANOVA with Dunnett’s multiple-comparisons test, comparing each mAb to the negative control (mu-MAb4) based on the main column effect averaged across all concentrations. Significance levels are indicated as *, p<0.05; **, p<0.01; *** p<0.001; ****, p<0.0001. Values with p > 0.05 are unmarked.

**Figure 6.**
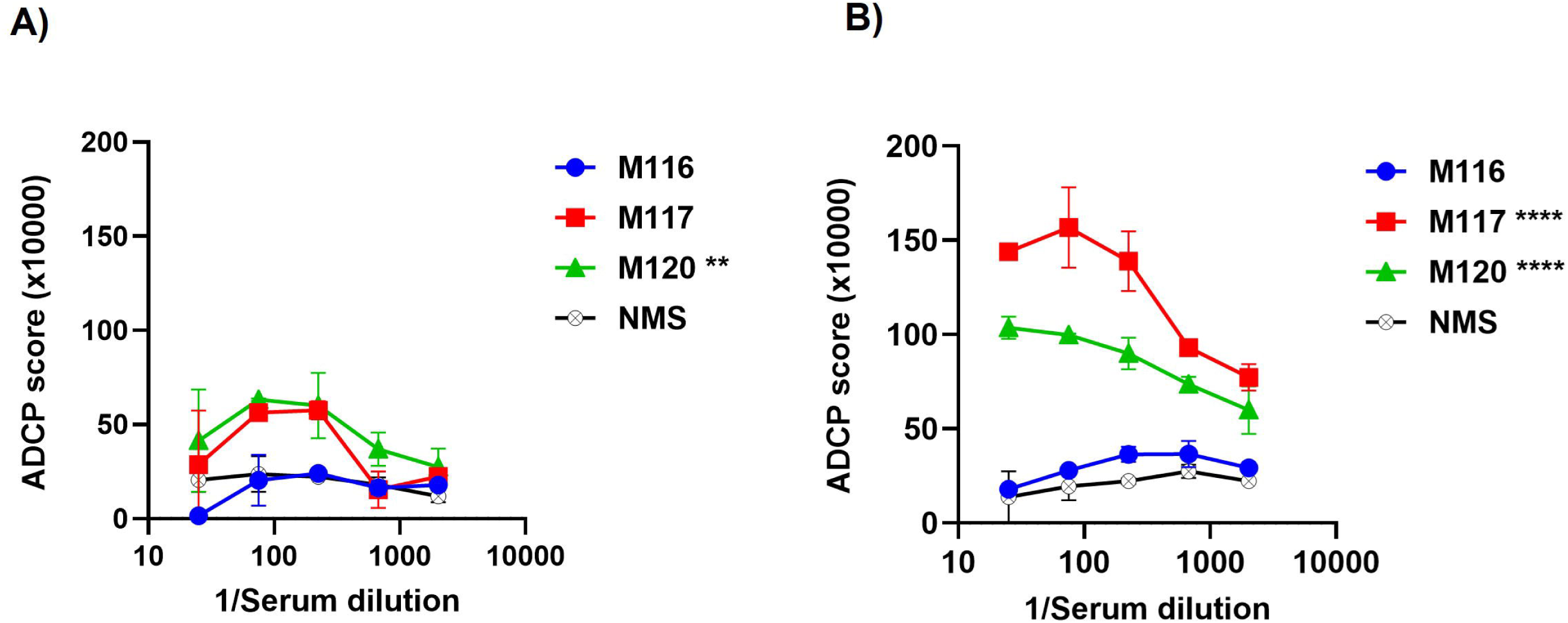
Antibody-dependent cellular phagocytosis (ADCP) mediated by sera from immunized mice against REJO HIV-1 virions and gp120. Sera from mice co-immunized with HIV-1 gp120 DNA + protein were tested for their ability to mediate ADCP using phagocytic THP-1 cells and fluorescent beads coated coupled to either AT-2 inactivated **(A)** REJO virions or **(B)** REJO gp120. ADCP scores are shown for individual vaccinated animals (M116, M117, M120) and normal mouse serum (NMS) as a negative control. Data represent the mean ± SD of replicate wells from one experiment. Statistical significance was determined using two-way ANOVA with Dunnett’s multiple-comparisons test comparing each vaccinated serum to NMS based on the main column effect averaged across dilutions (*P < 0.05; **P < 0.01; ***P < 0.001; ****P < 0.0001). Values with p > 0.05 are unmarked.

To evaluate the robustness of the virion-bead ADCP assay, we assessed both intra-assay precision and inter-assay reproducibility across three independent experiments performed on different days using the mouse monoclonal antibodies shown in Figure 5, each with RHPA virions derived from three independent virus preparations (Supplementary Table 1). Each antibody-virion condition was tested in duplicate wells in every run.

Intra-assay precision: Across the three experiments, conditions yielding measurable phagocytic activity showed low to moderate variability, consistent with expectations for flow-based cell assays. For antibodies with quantifiable signals, intra-assay coefficients of variation (CV%) generally ranged from 2.2% to 15.9%, with representative examples include 1B5 (CV 2.8-10.5%), 3F6 (CV 3.3-73.7%, signal-dependent), and 6C4 (CV 6.2-32.3%). As expected for flow-based cell assays, CVs increased substantially when mean activity approached background levels, including the Fc-impaired 1B5-LALA variant and the negative control antibody. Overall, these data demonstrate strong within-run repeatability for biologically meaningful signal ranges.

Inter-assay reproducibility: To assess run-to-run consistency mean ADCP scores for each antibody were compared across the three independent experiments. Inter-assay CVs ranged from 45.7% to 79.0% for antibodies exhibiting measurable activity, reflecting expected day-to-day variability associated with independent virus preparations and assay conditions. Importantly, despite variation in absolute signal magnitude, the relative ranking of antibody activities was highly preserved across experiments, indicating reproducible discrimination of functional differences among antibodies.

Together, these analyses demonstrate that the virion-bead ADCP assay exhibits high intra-assay precision and reliable inter-assay reproducibility for relative potency ranking, across independent experiments using different virion preparations. This performance profile is consistent with other complex, cell-based Fc-effector assays, where absolute signal intensity may vary between experiments, but functional ordering of antibodies is maintained.

## Discussion

Fc-mediated effector functions of antibodies, including ADCP, are increasingly recognized as important contributors to protective immunity against HIV-1, complementing Fab-mediated neutralization. Multiple studies in nonhuman primates have associated ADCP activity with reduced risk of infection (19, 20). In human vaccine trials such as RV144 and HVTN 505 ADCP was detected as part of the Fc mediated antibody response (21), highlighting the need for assay platforms that can accurately capture this function in a biologically meaningful context. Most ADCP studies rely on bead-based assays using recombinant Env proteins as phagocytic targets. These assays offer scalability and reproducibility, making them well suited for large comparative studies, however, they fail to replicate the membrane anchoring, structural organization and glycan complexity of the native Env trimer present on virions, features that critically influence antibody binding and Fc-mediated effector function in vivo (22, 23, 31). In contrast, infected cell-based assays that provide more physiologically relevant antigen presentation but are technically demanding and exhibit variability in antigen density limiting standardization and broader applications (28). Together, these limitations highlight the need for an assay that preserves native Env architecture on intact virions while remaining scalable and reproducible.

Native-like SOSIP Env trimers have been widely adopted as soluble antigens for ADCP and other Fc effector function assessments because they preserve many structural features of the native Env spike. Martin et al used biotinylated BG505 SOSIP trimers immobilized on beads and THP-1 effector cells to demonstrate robust ADCP triggered by rhesus macaque mAb (61). In clinical vaccine studies, serum antibodies elicited by ConM SOSIP.v7 immunization mediated ADCP of SOSIP-conjugated beads, supporting the use of SOSIP trimers to quantify Fc effector responses across species (62). Similarly, analyses of polyclonal IgG from HIV-infected individuals showed that binding to native-like trimeric SOSIPs correlated with phagocytic activity in bead-based formats (63). Despite these advantages, SOSIP trimers lack the membrane context and native lipid environment of virion-associated Env and are truncated constructs that do not contain the full gp41 ectodomain and membrane-proximal regions. These differences can influence epitope accessibility, glycan heterogeneity, and Fcγ receptor engagement. Moreover, not all HIV-1 isolates can be readily engineered or stably produced as SOSIP trimers, restricting their use to a limited subset of strains. This constraint is particularly relevant given that the same antibody can exhibit pronounced strain-specific differences in Fc effector activity, as observed in our virion-based assay, underscoring the importance of evaluating antibody function across multiple viral strains. Thus, while SOSIP-based assays are well suited for standardized and high-throughput screening, virion-based platforms provide complementary insight into Fc-mediated antibody function under structural constraints that more closely approximate intact HIV-1 particles.

Several prior studies have provided important insights into Fc-mediated interactions between antibodies and HIV. Snow et al. demonstrated that bNAbs mediate efficient ADCP of HIV-infected cells, whereas non-neutralizing antibodies (nnAbs) do not, highlighting the importance of native Env expression on cellular membranes for Fc-mediated clearance (31). Notably, nnAbs induced by the RV144 vaccine regimen have shown to mediate ADCP and were key correlates of protection (64). However, this infected-cell based format does not directly assess antibody-mediated clearance of cell-free virions and is inherently less amenable to standardization and high-throughput application. In addition, studies have shown that uncleaved form of Env is transported to the cell surface, but this nonfunctional Env is mostly excluded from the virus (65), thus while the assay present Env in relevant trimeric conformation it does not recapitulate the Env on virion surface completely. Tay et al. examined antibody-dependent internalization of infectious HIV-1 virions and showed that isotype and subclass influence virion uptake, providing critical insight into Fc-Fcγ receptor interactions (23). While informative, this approach relied on infectious virus and spinoculation, introducing technical constraints and limiting scalability.

The virion-based ADCP assay described here builds upon these conceptual advances by combining preserved, membrane-embedded Env presentation with a standardized and scalable bead-based format. In this assay, inactivated virions are covalently coupled to fluorescent beads and used as phagocytic targets. Although the beads are larger than HIV virions, the objective is not to model virion size but to preserve the native molecular organization of Env that governs antibody binding and Fcγ receptor engagement. This design is conceptually aligned with prior work emphasizing the importance of target architecture and Fc-Fcγ receptor interactions in determining phagocytic outcomes (23). By leveraging bead coupling as a phagocytosis-competent scaffold while preserving native Env trimer organization and glycan shielding, the assay enables reproducible, high-throughput measurement of antibody-mediated phagocytosis without the biosafety and variability concerns associated with free virions or infectious virus assays.

Our findings differ from those reported by Gach et al., who concluded that HIV-1 virions are largely resistant to ADCP due to insufficient surface Env density (32). This inefficiency was attributed to the low density of Env trimers on the virion surface, which limits effective Fcγ receptor engagement. In contrast, we and others, including Tay et al., observed measurable ADCP of HIV-1 virions mediated by diverse monoclonal antibodies and polyclonal sera. In our study, virions equivalent to ∼175 ng Env were used per coupling reaction, corresponding to a theoretically estimated average of ∼one virion per bead, a configuration unlikely to artificially increase antigen density relative to free virions (see Methods). We propose that the bead-coupling strategy enhances spatial organization and consistency of Env presentation, facilitating productive Fcγ receptor engagement without altering intrinsic virion properties.

Using this platform we observed clear strain- and epitope-dependent differences in ADCP activity across HIV-1 isolates. We observed marked differences in ADCP activity between CMU06 and REJO virions for several monoclonal antibodies, particularly the V3-directed mAb 2219, which was active against CMU06 but inactive against RHPA and REJO virions. These restrictions were not recapitulated when recombinant REJO gp120 was used as the target, where 2219 exhibited strong ADCP activity. This pattern is consistent with established models of Env structure in which the V3 loop is conformationally masked on native trimers by quaternary constraints and glycan shielding, limiting antibody access (52, 66–68). In contrast, monomeric gp120 lacks these quaternary constraints, leading to increased epitope exposure and functional readouts that may overestimate antibody activity relative to physiological conditions. Because antigen density and presentation differ fundamentally between virion- and protein-based targets, we did not perform direct quantitative comparisons of ADCP magnitude across assay formats. Instead, statistical analyses were conducted within each format relative to appropriate negative controls. While several antibodies showed statistically significant ADCP activity, the magnitude and consistency of these responses varied substantially. This highlights an important distinction between statistical significance and biological effect size, particularly in sensitive functional assays, and underscores the value of interpreting ADCP data in the context of antigen presentation and effect magnitude rather than p-values alone.

Importantly, the virion-based ADCP assay was effective across diverse experimental settings, detecting phagocytosis mediated by broadly neutralizing and non-neutralizing monoclonal antibodies, polyclonal sera from people living with HIV, and vaccine-elicited antibodies in mice. The assay was reproducible across independent virus preparations and compatible with murine antibodies, supporting its utility for preclinical vaccine evaluation. While THP-1 cells were used here as a standardized effector population to ensure scalability and reproducibility, future studies incorporating primary myeloid cells will further define the in vivo relevance of these findings.

In summary, the virion-based ADCP assay described here provides a practical and biologically relevant platform for measuring antibody-mediated phagocytosis of intact HIV-1 particles. By preserving key features of native Env while enabling standardized, high-throughput analysis, this approach complements existing ADCP assays and enables more nuanced evaluation of Fc effector function. Beyond HIV-1, the platform can be readily adapted to other enveloped viruses where native glycoprotein organization critically influences antibody activity and immune clearance.

## Author Contributions

CU conceived and designed the study. PR performed the experiments. CU and PR analyzed the data. CU wrote the original manuscript. CU and PR read and approved the final manuscript.

## Acknowledgments

The authors would like to thank Dr. Susan Zolla-Pazner and Ms. Xiaomei Liu for providing HIV-1-specific mAbs and Dr Andrew Duty for mouse mAb sequence.

## Data Availability Statement

The datasets generated for this study are available on request to the corresponding author.

